# Prediction of RNA-interacting residues in a protein using CNN and evolutionary profile

**DOI:** 10.1101/2022.06.03.494705

**Authors:** Sumeet Patiyal, Anjali Dhall, Khushboo Bajaj, Harshita Sahu, Gajendra P.S. Raghava

**Affiliations:** Department of Computational Biology, Indraprastha Institute of Information Technology, Okhla Phase 3, New Delhi-110020, India; Department of Computer Science and Engineering, Indraprastha Institute of Information Technology, Okhla Phase 3, New Delhi-110020, India

**Keywords:** RNA-interacting residues, Binary profile, Evolutionary profile, Convolutional neural network, Machine learning techniques

## Abstract

This paper describes a method Pprint2, which is an improved version of Pprint developed for predicting RNA-interacting residues in a protein. Training and validation datasets used in this study comprises of 545 and 161 non-redundant RNA-binding proteins, respectively. All models were trained on training dataset and evaluated on the validation dataset. The preliminary analysis reveals that positively charged amino acids such as H, R, and K, are more prominent in the RNA-interacting residues. Initially, machine learning based models have been developed using binary profile and obtain maximum area under curve (AUC) 0.68 on validation dataset. The performance of this model improved significantly from AUC 0.68 to 0.76 when evolutionary profile is used instead of binary profile. The performance of our evolutionary profile based model improved further from AUC 0.76 to 0.82, when convolutional neural network has been used for developing model. Our final model based on convolutional neural network using evolutionary information achieved AUC 0.82 with MCC of 0.49 on the validation dataset. Our best model outperform existing methods when evaluated on the validation dataset. A user-friendly standalone software and web based server named “Pprint2” has been developed for predicting RNA-interacting residues (https://webs.iiitd.edu.in/raghava/pprint2 and https://github.com/raghavagps/pprint2)

**Key Points:**

- Machine learning based models were developed using different profiles
- PSSM profile of a protein was created to extract evolutionary information
- PSSM profiles of proteins were generated using PSI-BLAST
- Convolutional neural network based model was developed using PSSM profile
- Webserver, Python- and Perl-based standalone package, and GitHub is available

**Author’s Biography:**

1. Sumeet Patiyal is currently working as Ph.D. in Computational Biology from Department of Computational Biology, Indraprastha Institute of Information Technology, New Delhi, India.
2. Anjali Dhall is currently working as Ph.D. in Computational Biology from Department of Computational Biology, Indraprastha Institute of Information Technology, New Delhi, India.
3. Khushboo Bajaj is currently working as MTech in Computer Science and Engineering from Department of Computer Science and Engineering, Indraprastha Institute of Information Technology, New Delhi, India.
4. Harshita Sahu is currently working as MTech in Computer Science and Engineering from Department of Computer Science and Engineering, Indraprastha Institute of Information Technology, New Delhi, India.
5. Gajendra P. S. Raghava is currently working as Professor and Head of Department of Computational Biology, Indraprastha Institute of Information Technology, New Delhi, India.

## Introduction

Proteins and RNA are the most crucial biological components of life, whereas RNA forms the essential part of ribosome, spliceosome and performs diverse roles within cell [1]. The RNA-protein interactions are necessary for several biological functions such as gene expression regulation, viral assembly & replication, posttranscriptional modification, and protein synthesis [2-7]. Recent studies reveal that RNA-protein interactions shows major involvement in developing human cancers and neurological disorders such as amyotrophic lateral sclerosis and Alzheimer’s [8-12]. These interaction also play a very crucial role in various infectious and genetic disorders [13-15]. In order to understand the functions and mechanisms of any biological process, it is necessary to get information regarding the RNA-protein interaction residues. In addition to that, RNA-protein interactions play major role in cellular homeostasis and malfunctioning in these interactions may lead to abnormal cellular functions and diseases [16-19]. Therefore, identification of the RNA-interacting residues can help with biotechnological manipulation. With the better understanding of the RNA-protein interacting residues one can design the RNA-based therapy to treat several RNA associated disorders [20-23]. The advancements in experimental techniques such as X-ray crystallography and nuclear magnetic resonance (NMR) several structures of protein-RNA interacting residues have been discovered and reported in Protein Data Bank [24]. However these experimental methods are very cost expensive and time-consuming. Whereas, computational approaches based on sequence information are very viable and cost-efficient in the probable detection of RNA-binding residues. Identification of RNA interacting residues/sites in proteins is a highly significant and complex processes in molecular biology, and it has drawn many researchers to work on it for several years in order to provide the best of the best research in this field.

In the last few years a wide range of computational algorithms has been developed for the prediction and identification of RNA-interacting proteins and residues. Broadly, these methods can be divided into two categories, sequence-based and structure-based methods [25-29]. In case of structure based methods, RNA-interacting residues are identified from structure of RNA-protein complex. Unfortunately, due to limitation of experimental techniques it is not possible to determine structure of RNA binding protein. In the last two decades; numerous sequence based methods have been developed where RNA-interacting residues are identified from sequence only. Kumar et. al., [30] developed a sequence based model using machine learning techniques and evolutionary information. Song et.al, [31] developed two approaches for predicting RNA-protein interaction residues in protein sequences; first is sequence-based method and second is feature-based method. In addition, PredRBR integrated huge number of sequence, structure-based features for the prediction of protein-RNA binding affinity [32]. Another computational method named RPiRLS [33] predicts RNA-protein interactions using the structural information. Recently developed methods such as ProNA2020 and hybridNAP predicts using sequence information [34].

In this study we have made a systematic attempt to construct machine learning and deep learning based models using the sequence information. We compute physio-chemical properties, binary profile and position specific scoring matrix or evolutionary profile based features. In order to develop prediction models, we used a wide range of machine learning algorithms such as decision tree, extra-tree classifier, random forest, naive Bayes, gradient boosting, logistic regression, and k-nearest neighbour. In addition, models have been developed using deep learning particularly convolutional neural network. In order to provide uniform benchmarking, all existing methods have been evaluated on a validation dataset. We have integrated our best models in the webserver and standalone package for the prediction of RNA-interacting or non-interacting residues in a protein sequence. It has been shown that our convolutional neural network based model using evolutionary profile better than existing methods.

## Material and Methods

### Dataset Collection and Pre-processing

In the current study, we have compiled the datasets from the recently published paper HybridNAP [35] and ProNA2020 [34], which consist of 1057 and 360 annotated protein sequences, respectively. We have used CD-HIT software with the standards of 30% sequence identity to take care the redundancy, which resulted in the 545 protein sequences in the training dataset and 161 sequences in the validation dataset. Eventually, we left with 18559 RNA-interacting and 171879 non-interacting residues in the training dataset using 545 sequences. In case of validation dataset we obtained 6966 interacting and 44349 non-interacting residues using 161 sequences. After that, we generate the overlapping patterns for each sequence and with length 17, as done in previous studies [36]. The central i.e. 9th residue was taken as the representative of whole pattern, such as if the 9th residue is RNA-interacting the pattern is assigned as RNA-interacting otherwise non-interacting. To handle the terminal residues, eight “X” residues were added to both terminals of the sequences before generating the patterns, so that each residue gets the chance to be the central residue. Figure 1 represents the complete workflow adapted in this study.

**Figure 1:**
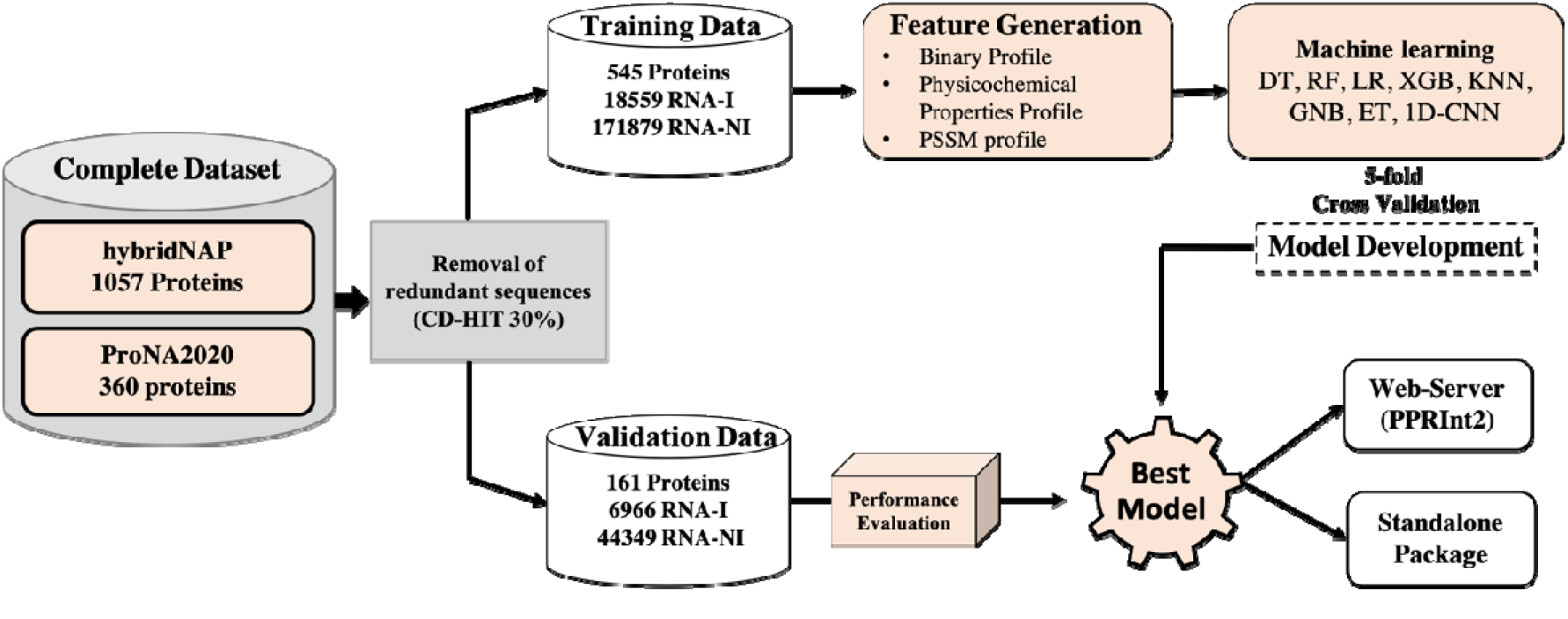
Complete workflow of the study including the dataset creation, feature generation, model development and evaluation.

### Amino acid composition

Amino acid composition (AAC) of residues in a protein sequence was computed using Pfeature standalone version [37]. For AAC, it calculate the percent composition and generate a fixed length vector of 20 amino acid residues as shown in equation 1. where is amino acid composition of residue type i; R_i_ and L number of residues of type i and length of sequence.

### Feature Generation

#### Binary Profile

The binary profile is generated with the help of Pfeature [37], here we computed binary profile. Where, each amino acid residue represented with a fixed vector size i.e. 21, for example A is represented as 1,0,0,0,0,0,0,0,0,0,0,0,0,0,0,0,0,0,0,0,0; in which 20 are natural amino acids and one is for dummy variable, whereas X is denoted as 0,0,0,0,0,0,0,0,0,0,0,0,0,0,0,0,0,0,0,0,1. Hence, each pattern is denoted by a fixed length vector of size 357 (17*21).

#### Physicochemical Properties Profile

We have implemented Pfeature [37] to calculate physicochemical properties profile using binary profile module. In this approach, each amino acid is represented with a vector of size 25, where each element denotes the particular physicochemical property, such as amino acid “A” is represented as 0,0,1,0,1,1,0,0,0,0,1,1,0,0,0,0,1,0,0,1,0,0,1,1,0; where “X” is denoted by zero vector of length 25. ‘1’ denotes the presence and ‘0’ denoted the absence of a particular property. Since, the size of each pattern is 17, hence the resulting length for each vector is 425 (17 * 25).

#### Evolutionary Profile

In order to compute the evolutionary information of residues we generate Position-Specific Scoring Matrix (PSSM) profiles. The PSSM profile was created using PSI-BLAST, which searches each sequence against the Swiss-Prot database. PSI-BLAST was conducted with three iterations along with an e-value of 1e^-3^. As shown in equation 2, we further normalized the profiles, where each pattern is represented as a vector of length 357 in the final matrix for each sequence, which is of dimension Nx21, where N is the length of the protein sequence.

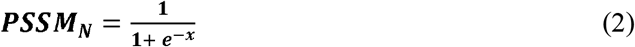

Where, x is the PSSM score and PSSMN is the normalized value.

### Model Building

In order to classify RNA-interacting and non-interaction residues in a protein sequence, we have implemented various classifier. These machine learning classifiers implemented in python library Scikit-learn [38]. To develop prediction models, we implement various classifiers such as Decision Tree (DT), Random Forest (RF), Logistic Regression (LR), eXtreme Gradient Boosting (XGB), Gaussian Naive Bayes (GNB), Extra-Tree Classifier (ET), K-nearest neighbor, and One-Dimensional Convolutional Neural Network (1D-CNN).

### Five-fold Cross Validation

To train, test, and validate the prediction models, we employed 5-fold cross-validation and an external validation as implemented in previous studies [39, 40]. 5-fold cross-validation was employed only on the training dataset. In which the data was split into five parts, where four of which were used to train the model and the fifth set was utilised for testing. This process is iterated five times, in which each set is used in training and testing the models.

### Performance Evaluation

In this work, we examined the performance of the model using sensitivity, specificity, f1 score, accuracy, area under the receiver operating characteristic (AUROC), and Matthews correlation coefficient (MCC). These parameters belong to threshold dependent (i.e., sensitivity, specificity, accuracy and MCC) and independent category (AUROC) to evaluate our models. These parameters were computed using the following equations (3-7).

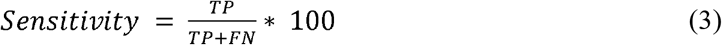

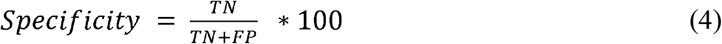

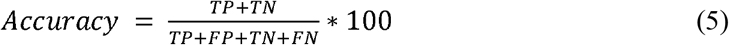

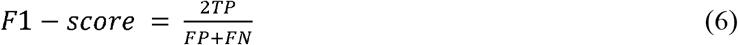

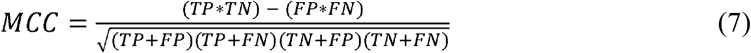

Where, FP is false positive, FN is false negative, TP is true positive and TN is true negative.

## Results

### Compositional Analysis

In order to understand the distribution of amino acid residues in the RNA-interacting vs non-interacting vs general proteome, we have calculated the percent amino acid composition. For percent amino acid composition of general proteome, we have used the values from the Swiss-Prot database available at https://web.expasy.org/protscale/pscale/A.A.Swiss-Prot.html. As exhibited by Figure 2, positively charged residue H, K, and R are abundant in RNA-interacting residues as compare to non-interacting or general proteome, where A, D, E, I, L, and V are rich in non-interacting and general proteome.

**Figure 2:**
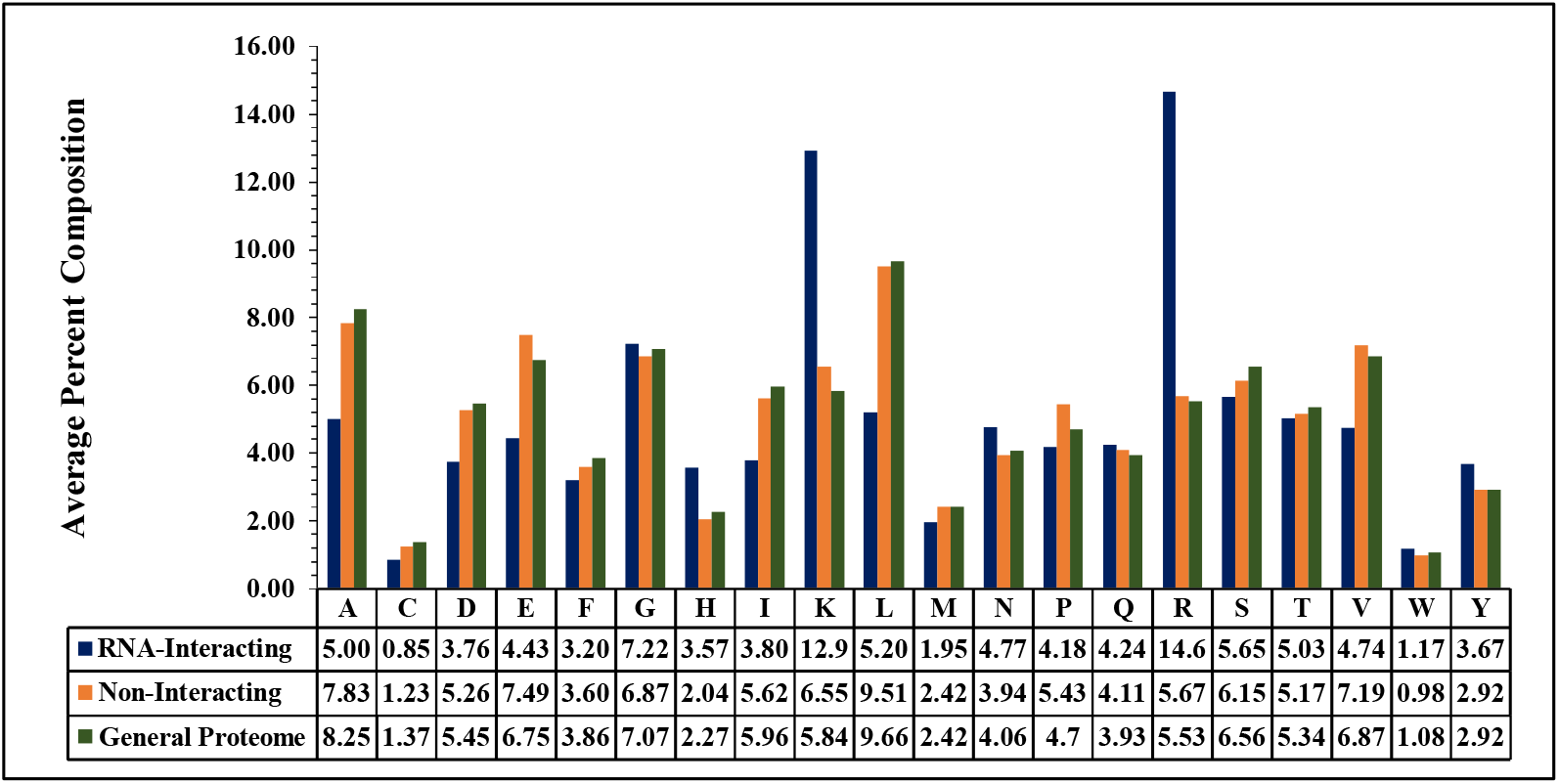
Percent composition of each residue in RNA-interacting, non-interacting residues, and general proteome.

### Physiochemical Property Analysis

In order to explore the nature of amino acids involved in the RNA-interaction, we computed physiochemical properties-based percent composition of residues involved in RNA-interaction. As shown by Figure 3, positively charged, basic and hydrophilic residues are abundant in the RNA-binding sites as RNA has a negative backbone. On the other hand, negatively charged, neutral charged, acidic, hydrophobic, small, non-polar, and aliphatic residues are scare in RNA-interacting sites.

**Figure 3:**
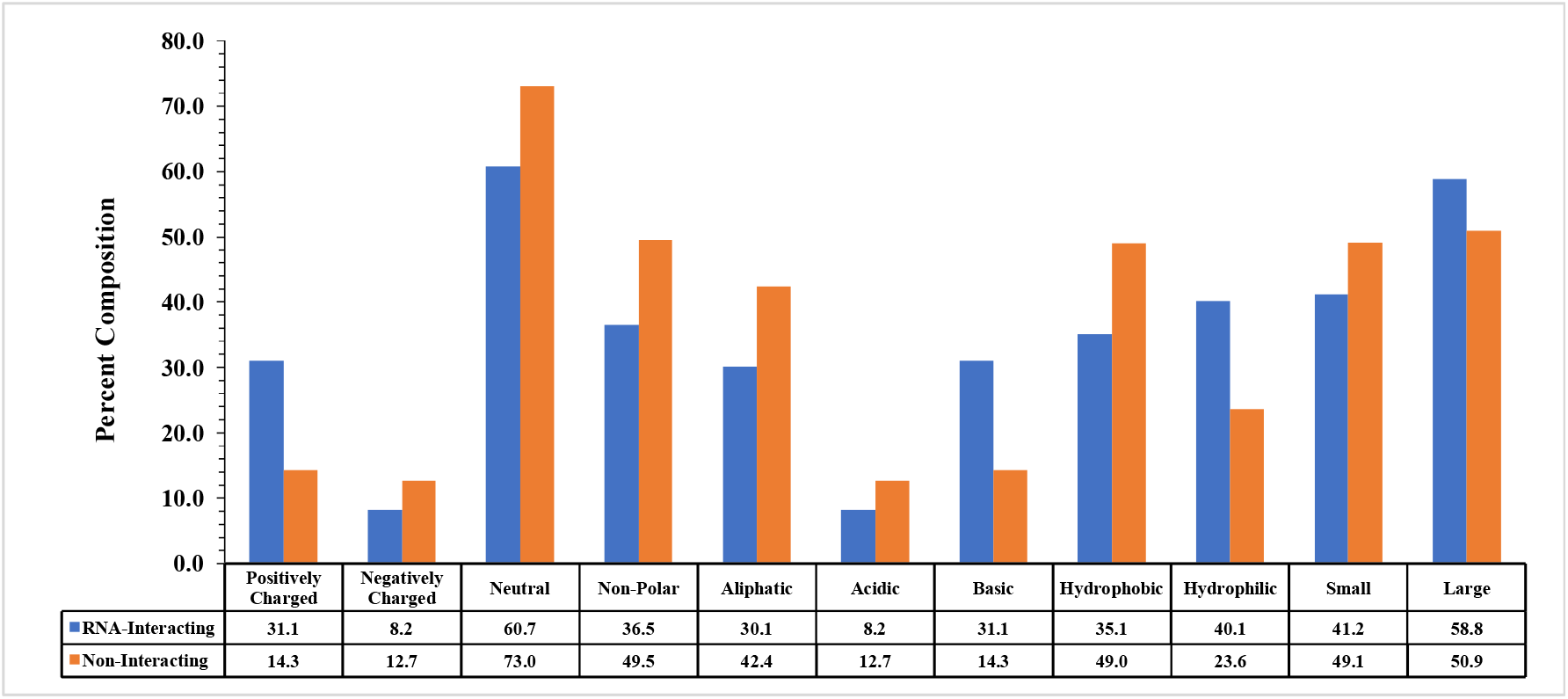
Percent composition based on physiochemical properties of residues involved in RNA-interaction.

### Two Sample Logo

In the past, number of studies have shown that the interaction property of a residue is influenced by its neighbouring residues. In this study, we have created the patterns of length 17, where central i.e. 9^th^ residue is the representative of whole pattern and based on that a pattern is assigned as interacting or non-interacting. Figure 4 represents the preference of residues as each position in RNA-interacting and non-interacting patterns, and it shows that residue R is highly preferred at interacting site followed by K and H, flanked by positively charged residues R and K. On the contrary, non-interacting sites are preferred by residue L, followed by E, A and V, where these residues are flanked by residues E and L.

**Figure 4:**
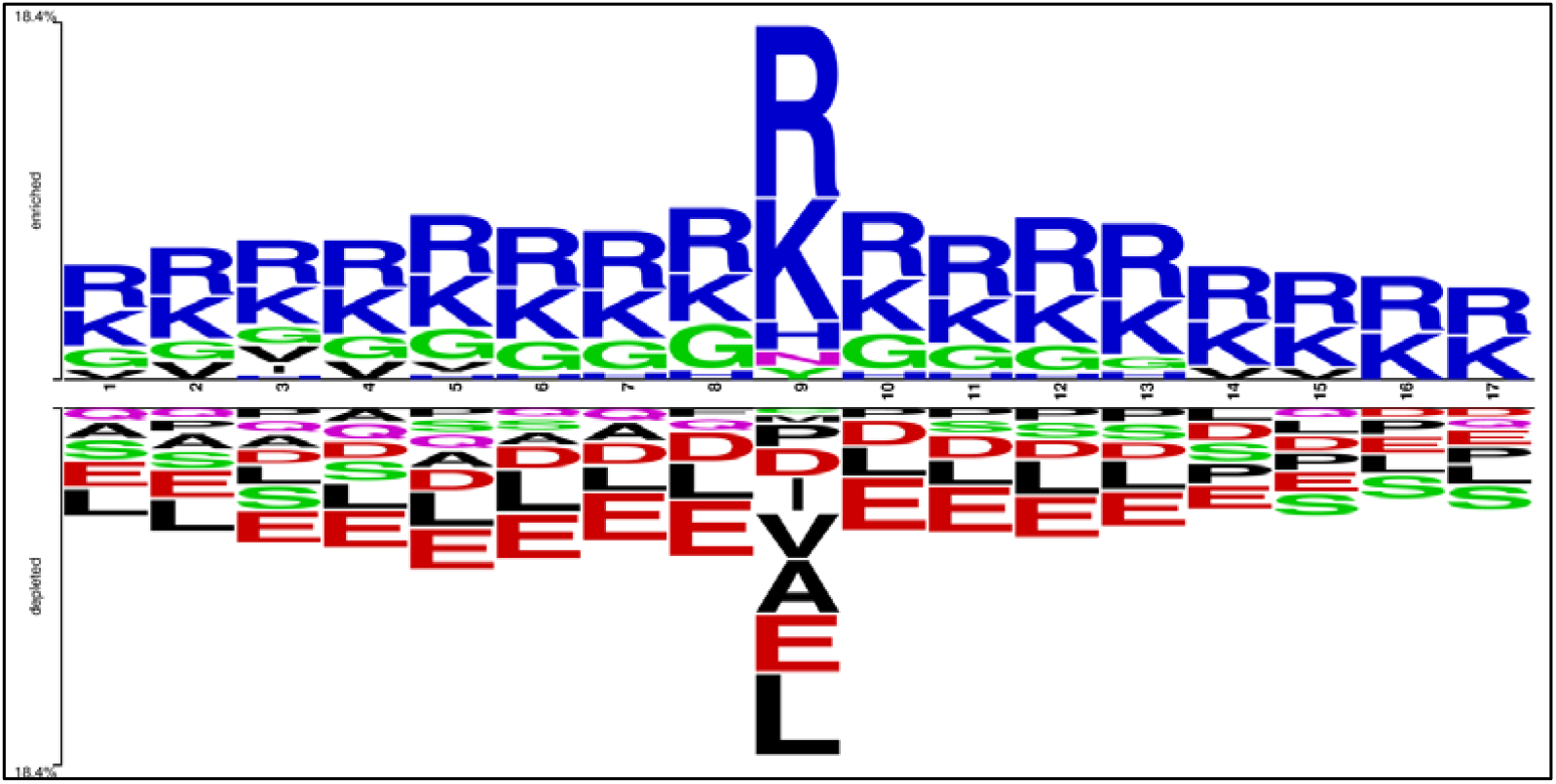
Two sample logo exhibits the preference of residues at each position in RNA-interacting and non-interacting patterns.

### Performance of Binary Profile

We have used binary profile as input features to train and evaluate the prediction models by implementing various machine learning classifiers. We used five-fold cross validation technique to build prediction models on training datasets. As shown in Table 1, in case of machine learning techniques both logistic regression and XGB achieve maximum AUC 0.74. In case of deep learning, 1D-CNN based model obtained AUC 0.81, which is highest. In order obtain unbiased performance of models we evaluate these models on validation dataset. In case of machine learning techniques, we achieved maximum AUC 0.68 for both logistic regression and XGB based models. In case of 1D-CNN performance of model decrease drastically from 0.81 to 0.68 when tested on validation dataset instead of training dataset. In summary, both 1D-CNN and machine learning based models achieve same performance on validation dataset.

**Table 1:** The performance binary profile based on models developed using different classifiers.

| Classifier | Training Dataset |  |  |  |  |  |  | Validation Dataset |  |  |  |  |  |  |
| --- | --- | --- | --- | --- | --- | --- | --- | --- | --- | --- | --- | --- | --- | --- |
|  | Sens | Spec | Acc | AUC | F1 | K | MCC | Sens | Spec | Acc | AUC | F1 | K | MCC |
| DT | 19.14 | 88.93 | 82.13 | 0.54 | 0.17 | 0.07 | 0.07 | 15.94 | 89.07 | 79.14 | 0.53 | 0.17 | 0.05 | 0.05 |
| RF | 66.57 | 67.53 | 67.43 | 0.73 | 0.29 | 0.16 | 0.21 | 55.77 | 68.54 | 66.81 | 0.67 | 0.31 | 0.15 | 0.18 |
| LR | 67.20 | 68.16 | 68.07 | 0.74 | 0.29 | 0.16 | 0.22 | 55.79 | 70.37 | 68.39 | 0.68 | 0.32 | 0.16 | 0.19 |
| XGB | 66.88 | 67.54 | 67.48 | 0.74 | 0.29 | 0.16 | 0.21 | 55.86 | 69.65 | 67.78 | 0.68 | 0.32 | 0.16 | 0.19 |
| KNN | 60.32 | 63.75 | 63.41 | 0.64 | 0.24 | 0.10 | 0.15 | 52.60 | 64.66 | 63.02 | 0.60 | 0.28 | 0.10 | 0.12 |
| GNB | 66.63 | 65.14 | 65.28 | 0.71 | 0.27 | 0.14 | 0.19 | 56.30 | 67.11 | 65.64 | 0.65 | 0.31 | 0.14 | 0.17 |
| ET | 68.25 | 65.72 | 65.97 | 0.73 | 0.28 | 0.15 | 0.21 | 58.38 | 66.64 | 65.52 | 0.67 | 0.32 | 0.15 | 0.18 |
| 1D-CNN | 73.91 | 73.76 | 73.78 | 0.81 | 0.36 | 0.24 | 0.31 | 50.66 | 74.63 | 71.38 | 0.68 | 0.33 | 0.17 | 0.19 |

\* DT: Decision Tree; RF: Random Forest; LR: Logistic Regression; XGB: eXtreme Gradient Boosting; KNN: K-Nearest Neighbour; GNB: Gaussian Naïve Bayes; ET: Extra Tree; 1D-CNN: One-Dimensional Convolutional Neural Network; Sens: Sensitivity; Spec: Specificity; Acc: Accuracy; AUC: Area Under the receiver operating characteristic Curve; K: Kappa; MCC: Matthews Correlation Coefficient

### Performance of Physicochemical properties profile

In addition, models have been developed using physicochemical properties profile, which represents each amino acid with vector of length 25. In case of machine learning techniques, we got maximum AUC 0.74 and 0.68 on training and validation dataset respectively (Table 2).

**Table 2:** Performance measures for models developed by implementing various classifiers using physicochemical properties profile as the input feature.

| Classifier | Training Dataset |  |  |  |  |  |  | Validation Dataset |  |  |  |  |  |  |
| --- | --- | --- | --- | --- | --- | --- | --- | --- | --- | --- | --- | --- | --- | --- |
|  | Sens | Spec | Acc | AUC | F1 | K | MCC | Sens | Spec | Acc | AUC | F1 | K | MCC |
| DT | 16.15 | 91.03 | 83.73 | 0.54 | 0.16 | 0.07 | 0.07 | 14.00 | 91.46 | 80.95 | 0.53 | 0.17 | 0.06 | 0.06 |
| **RF** | 67.74 | 64.56 | 64.87 | 0.72 | 0.27 | 0.14 | 0.20 | 56.88 | 66.37 | 65.08 | 0.66 | 0.31 | 0.14 | 0.17 |
| **LR** | 67.30 | 68.19 | 68.10 | 0.74 | 0.29 | 0.16 | 0.22 | 55.70 | 70.48 | 68.47 | 0.68 | 0.32 | 0.16 | 0.19 |
| **XGB** | 65.41 | 68.60 | 68.29 | 0.73 | 0.29 | 0.16 | 0.21 | 54.22 | 70.50 | 68.29 | 0.68 | 0.32 | 0.16 | 0.18 |
| **KN** | 62.43 | 63.32 | 63.23 | 0.65 | 0.25 | 0.11 | 0.16 | 53.22 | 63.81 | 62.37 | 0.60 | 0.28 | 0.10 | 0.12 |
| **GNB** | 65.44 | 66.15 | 66.08 | 0.71 | 0.27 | 0.14 | 0.19 | 54.19 | 68.58 | 66.63 | 0.66 | 0.31 | 0.14 | 0.16 |
| **ET** | 65.81 | 67.25 | 67.11 | 0.72 | 0.28 | 0.15 | 0.20 | 54.65 | 68.74 | 66.83 | 0.66 | 0.31 | 0.14 | 0.17 |
| **1D-CNN** | 69.84 | 69.60 | 69.62 | 0.77 | 0.31 | 0.19 | 0.25 | 56.42 | 71.14 | 69.14 | 0.69 | 0.33 | 0.17 | 0.20 |

\* DT: Decision Tree; RF: Random Forest; LR: Logistic Regression; XGB: eXtreme Gradient Boosting; KNN: K-Nearest Neighbour; GNB: Gaussian Naïve Bayes; ET: Extra Tree; 1D-CNN: One-Dimensional Convolutional Neural Network; Sens: Sensitivity; Spec: Specificity; Acc: Accuracy; AUC: Area Under the receiver operating characteristic Curve; K: Kappa; MCC: Matthews Correlation Coefficient

In case of deep learning 1D-CNN, we achieved maximum AUC 0.79 and 0.68 on training and validation dataset respectively. These results indicate that models based on physicochemical properties and binary profile have nearly same precision.

### Performance of Evolutionary Profile

It has been shown in the previous studies that evolutionary information provides more information then single sequence. Thus, in this study PSSM profiles were generated for proteins to capture evolutionary information. Machine learning based models have been developed using PSSM profile for predicting RNA interacting residues. As shown in Table 3, almost all classifies developed using PSSM profile perform better then binary profile-based models. In case of machine learning, XGB based model obtain AUC 0.82 and 0.76 on training and validation dataset respectively. It means the performance increases from 0.68 to 076 on validation dataset when we used PSSM profile in place of binary profile. It shows importance of PSSM profile in predicting RNA-interacting residues. Our 1D-CNN based model obtained AUC 0.91 and 0.82 on training and validation datasets respectively. In summary, 1D-CNN achieved maximum AUC 0.82 on validation dataset.

**Table 3:**
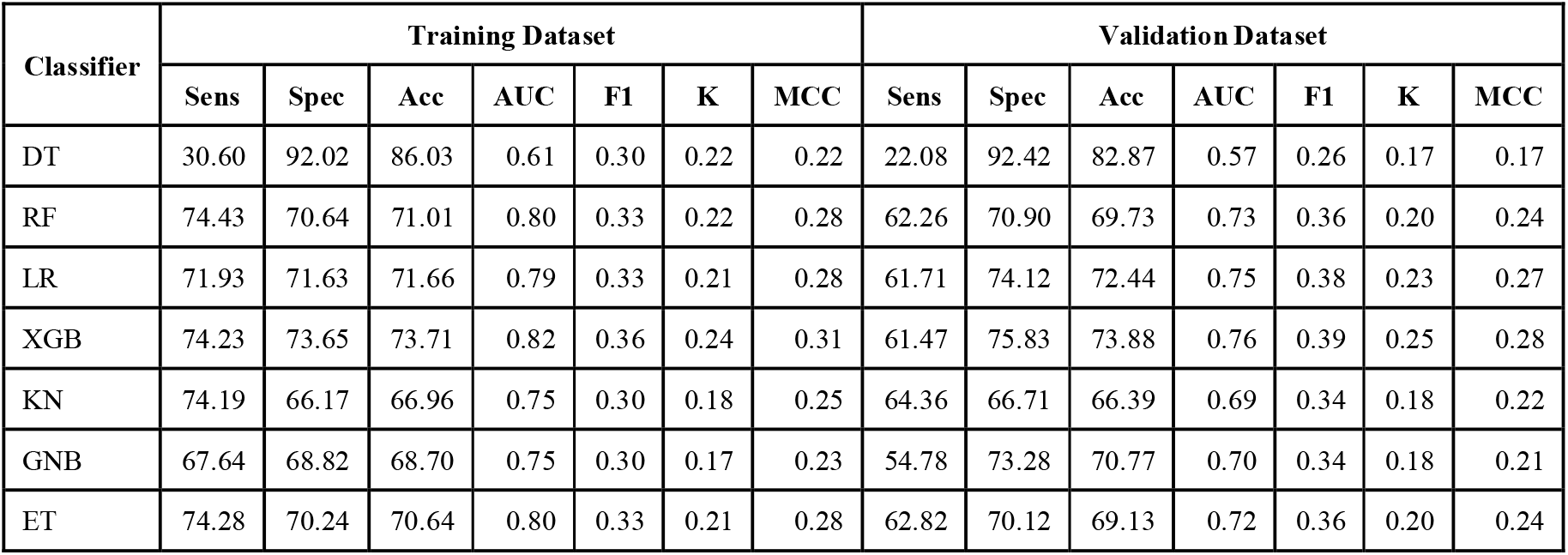
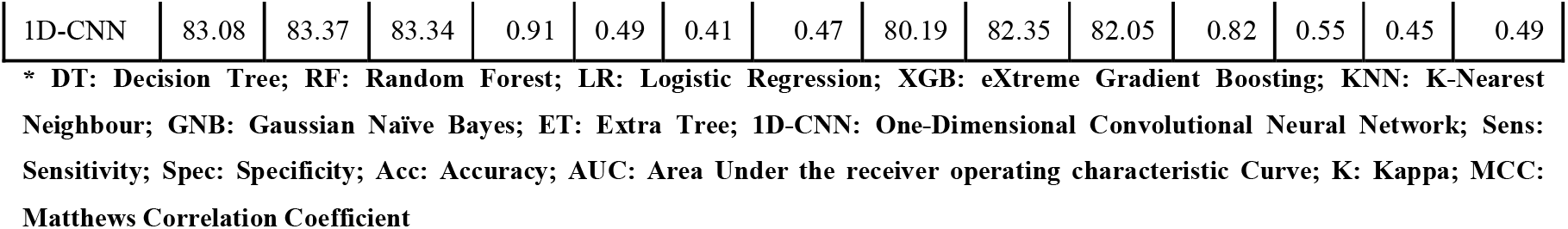
Performance of various classifiers using PSSM profile as input feature for training and validation dataset.

| Classifier | Training Dataset |  |  |  |  |  |  | Validation Dataset |  |  |  |  |  |  |
| --- | --- | --- | --- | --- | --- | --- | --- | --- | --- | --- | --- | --- | --- | --- |
|  | Sens | Spec | Acc | AUC | F1 | K | MCC | Sens | Spec | Acc | AUC | F1 | K | MCC |
| DT | 30.60 | 92.02 | 86.03 | 0.61 | 0.30 | 0.22 | 0.22 | 22.08 | 92.42 | 82.87 | 0.57 | 0.26 | 0.17 | 0.17 |
| RF | 74.43 | 70.64 | 71.01 | 0.80 | 0.33 | 0.22 | 0.28 | 62.26 | 70.90 | 69.73 | 0.73 | 0.36 | 0.20 | 0.24 |
| LR | 71.93 | 71.63 | 71.66 | 0.79 | 0.33 | 0.21 | 0.28 | 61.71 | 74.12 | 72.44 | 0.75 | 0.38 | 0.23 | 0.27 |
| XGB | 74.23 | 73.65 | 73.71 | 0.82 | 0.36 | 0.24 | 0.31 | 61.47 | 75.83 | 73.88 | 0.76 | 0.39 | 0.25 | 0.28 |
| KN | 74.19 | 66.17 | 66.96 | 0.75 | 0.30 | 0.18 | 0.25 | 64.36 | 66.71 | 66.39 | 0.69 | 0.34 | 0.18 | 0.22 |
| GNB | 67.64 | 68.82 | 68.70 | 0.75 | 0.30 | 0.17 | 0.23 | 54.78 | 73.28 | 70.77 | 0.70 | 0.34 | 0.18 | 0.21 |
| ET | 74.28 | 70.24 | 70.64 | 0.80 | 0.33 | 0.21 | 0.28 | 62.82 | 70.12 | 69.13 | 0.72 | 0.36 | 0.20 | 0.24 |
| 1D-CNN | 83.08 | 83.37 | 83.34 | 0.91 | 0.49 | 0.41 | 0.47 | 80.19 | 82.35 | 82.05 | 0.82 | 0.55 | 0.45 | 0.49 |

\* DT: Decision Tree; RF: Random Forest; LR: Logistic Regression; XGB: eXtreme Gradient Boosting; KNN: K-Nearest Neighbour; GNB: Gaussian Naïve Bayes; ET: Extra Tree; 1D-CNN: One-Dimensional Convolutional Neural Network; Sens: Sensitivity; Spec: Specificity; Acc: Accuracy; AUC: Area Under the receiver operating characteristic Curve; K: Kappa; MCC: Matthews Correlation Coefficient

**Table 5:** Performance of existing methods on the validation dataset.

| Methods | Sensitivity | Specificity | Accuracy | AUC | F1 | K | MCC |
| --- | --- | --- | --- | --- | --- | --- | --- |
| DRNAPred | 45.40 | 51.87 | 51.59 | 0.52 | 0.08 | 0.00 | 0.01 |
| HybridNAP | 56.90 | 56.05 | 56.09 | 0.59 | 0.10 | 0.02 | 0.05 |
| Pprint | 60.15 | 60.29 | 60.28 | 0.63 | 0.12 | 0.04 | 0.08 |
| ProNA2020 | 62.07 | 62.44 | 62.43 | 0.68 | 0.13 | 0.05 | 0.10 |
| RNABindRPlus | 67.24 | 68.36 | 68.31 | 0.73 | 0.16 | 0.09 | 0.15 |
| Pprint2 | 80.19 | 82.34 | 82.05 | 0.82 | 0.55 | 0.45 | 0.49 |

We have also compared the difference between the AUC of models built using PSSM profile, to understand the overfitting in the models by comparing their performances in training and validation dataset as shown in Figure 5. The difference in the AUC for training and validation dataset showed that DT model is least overfitted, whereas the performance of the deep learning model exhibits that the resulting model is quite overfitted as compare to the other models.

**Figure 5:**
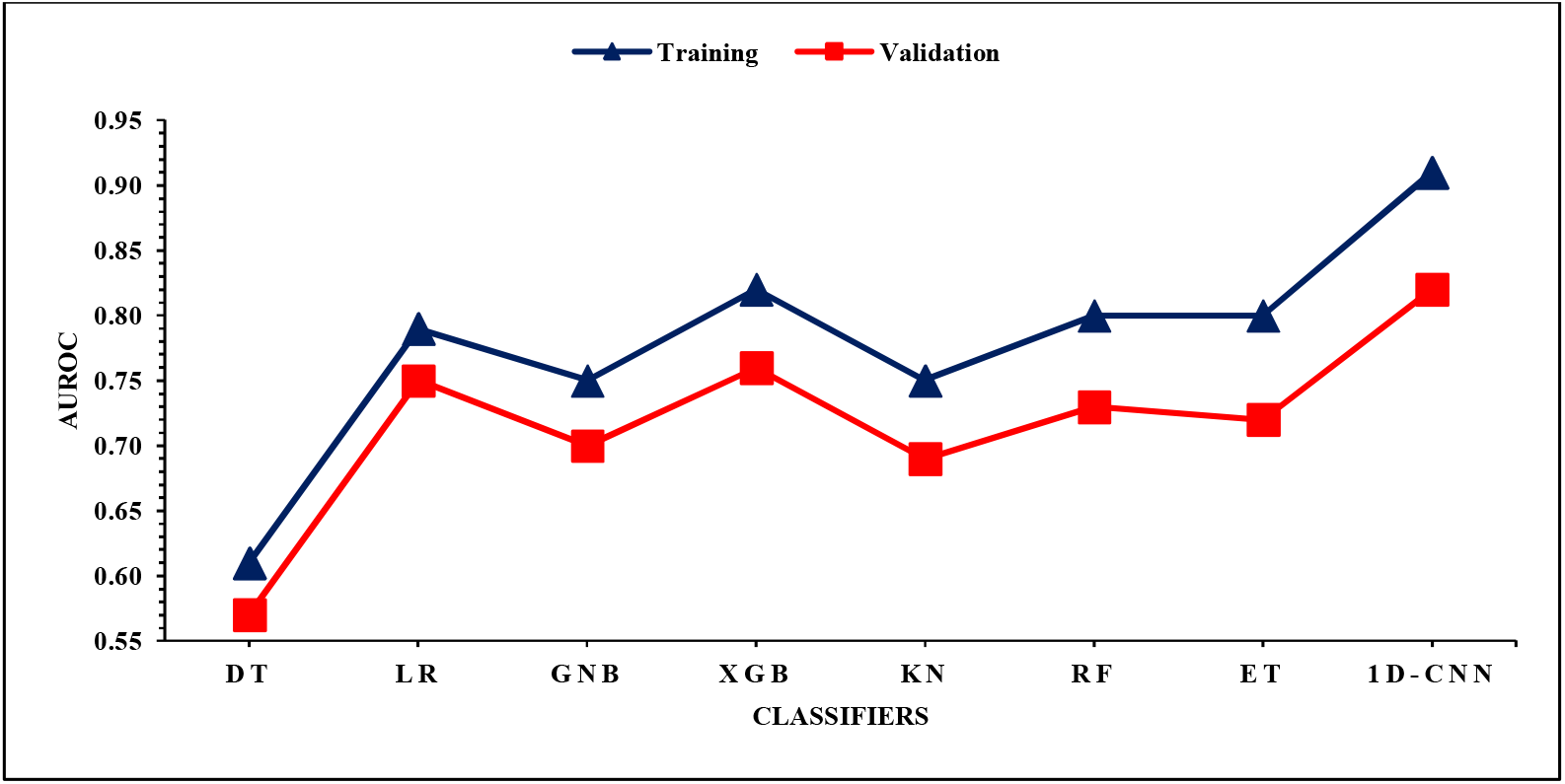
Difference in the AUC of various models developed using PSSM profile.

**Figure 6:**
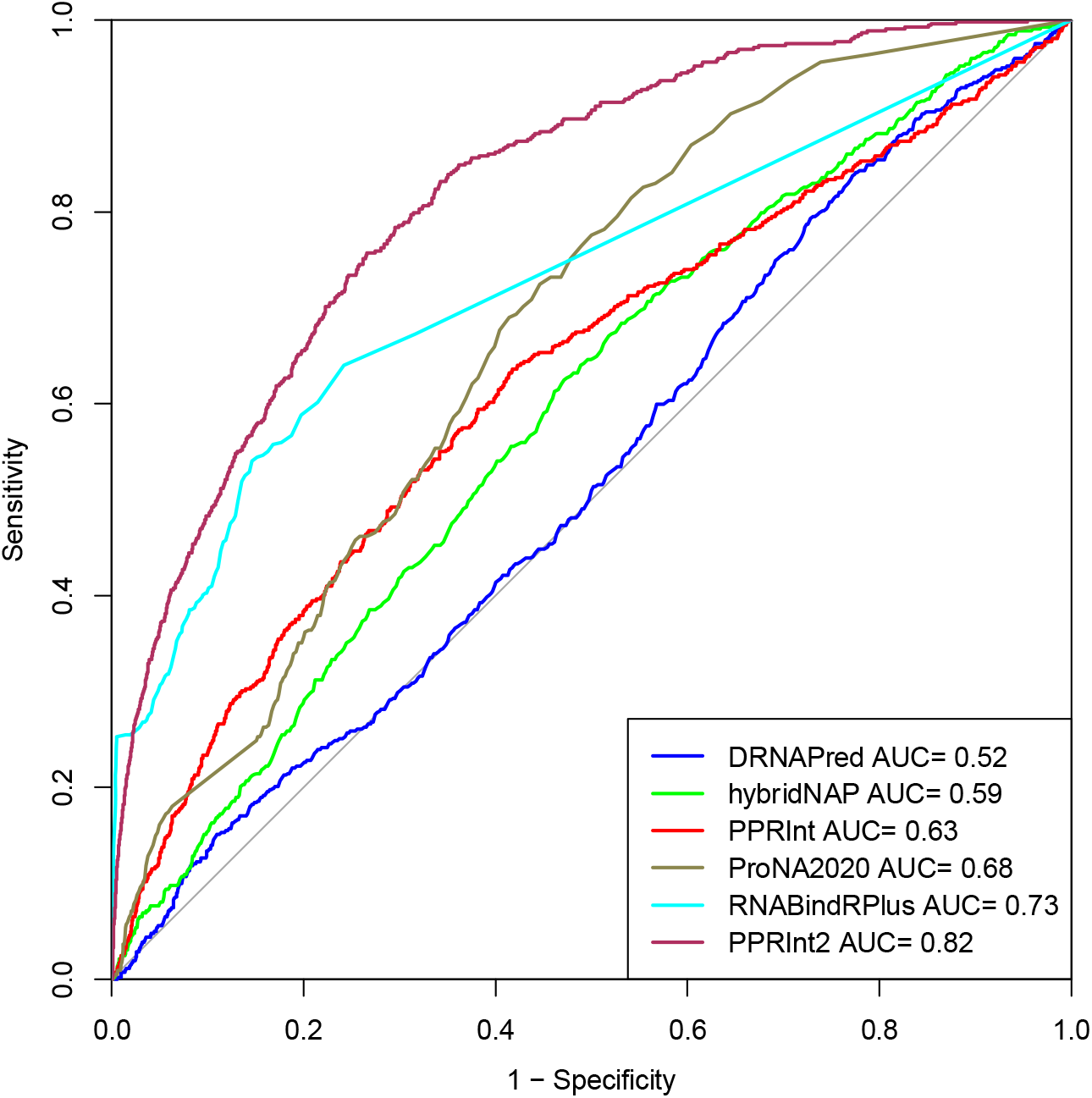
ROC curve shows the performance of all methods on validation dataset in term of area under curve (AUC)

### Comparison with Existing Methods

In order to justify a newly developed method, it is essential to compare its performance with the existing methods. The existing methods have been trained and evaluated on different datasets as these methods have been developed over the years. In order to provide unbiased evaluation of methods, one should evaluate performance of all methods on a common dataset. In this paper, evaluate the performance of the existing methods on validation dataset. The performance measures taken into consideration were sensitivity, specificity, AUC, accuracy, F1, Kappa, and MCC. As shown in Table 6, Pprint2 perform better than existing methods with AUC of 0.82, whereas RNABindRPlus performed second best among the existing methods with AUC 0.73.

### Service to Scientific Community

In order to facilitate scientific community, we developed a web server and standalone package called Pprint2, which is available at https://webs.iiitd.edu.in/raghava/pprint2/. Our web server allows the users to predict RNA interacting residues in a protein sequence pasted/uploaded by user. It allows users to select any modules for prediction that include PSSM profile and binary profile. The resulting page displays the input sequence(s) with highlighting the interacting residues in red colour as represented in Figure 7. In addition, server provide the facility to download the results in .txt, .pdf, and .png format. In addition, we have also provided the python-based standalone package, which is available from https://github.com/raghavagps/pprint2.

**Figure 7:**
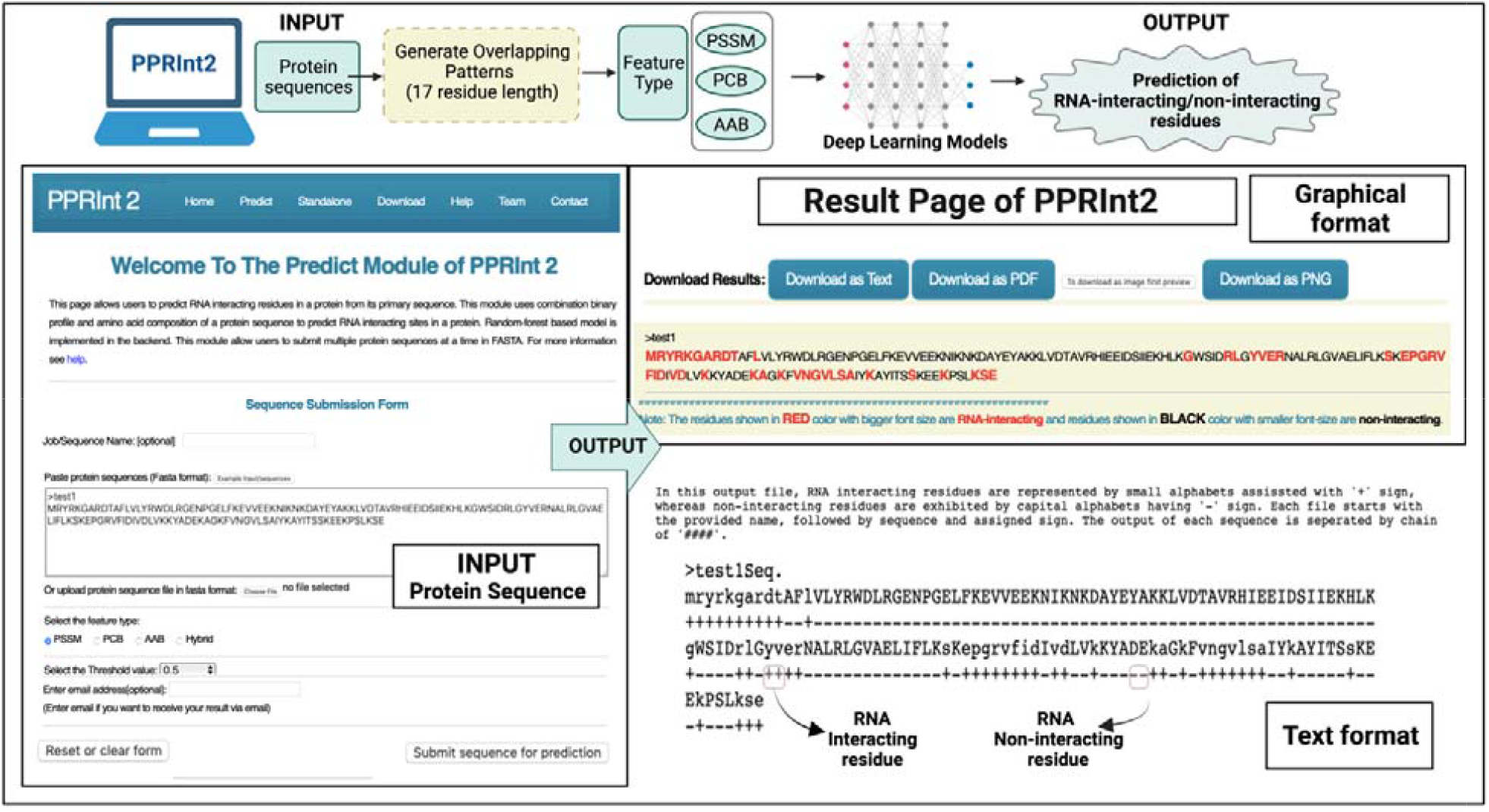
Complete usage of prediction module of webserver.

## Discussion and Conclusion

The interaction between RNA and protein complexes is responsible for many fundamental processes such as splicing, translation, transport, and silencing [41, 42]. Accurate identification of amino acid residues involving in the interaction with the RNA lead to the better understanding of the sequence specific mechanism of RNA-protein interaction [43, 44]. Correct identification of RNA-interacting residues in a protein is only possible if structure of RNA-protein complexes is available. Unfortunately, crystallization of all RNA-protein complexes is not possible due to number of limitation of experimental techniques like crystallography and NMR. In addition, experimental techniques are costly and time consuming. In order to facilitate researcher in the field of RNA biology, many methods have been developed for predicting RNA interacting residues [45-47]. One of the major limitation of previous methods was that they have been trained and evaluated on limited RNA binding proteins. For example, in our previous method Pprint [30], we trained and test our models on only 86 RNA binding proteins. Due to lack of sufficient data, a lenient cut-off level 70% has been used for removing redundant proteins. This is true for most of old methods where limited set of RNA binding proteins were used for training and evaluation. In addition, proteins in dataset contain highs level of similarity with each other. Over the years structures of RNA-protein complexes have been grown drastically in PDB [44, 48]. Thus, there is a need to develop new method on large set of RNA binding proteins whose structure is available in PDB. In addition, there is a need to create separate dataset for training and validation. In order to avoid any over optimization of machine learning models that proteins in training and validation should not have minimum similarity. In this study, we make an attempt to train our models on large set of proteins. Initially, we obtained 1057 and 360 protein sequences from recently published articles hybridNAP [49] and proNA2020 [34]. In order to create dataset of redundant RNA-binding protein, we removing the redundant sequences using CD-HIT at 30%. This lead to a dataset of 545 proteins in training and 161 sequences in validation dataset. This is one of the largest dataset used for training and validation. In addition, proteins in training and validation dataset have similarity 30% or less.

As shown in results section, in most of the cases performance of our machine learning based model on training and validation dataset is nearly same. This indicate that our models are not over optimized on training dataset. Our results indicate that performance of models improved significantly using evolutionary information. This support previous studies including our previous method Pprint that evolutionary information is important for prediction. The performance of models based on binary profile and physiochemical property is nearly same. It means over optimization is not a major challenge in case of machine learning techniques if models developed using cross-validation techniques. In case of 1D-CNN (deep learning), we observed high level of over optimization. Most of the 1D-CNN models have high performance on training dataset compare to training dataset. Thus cross-validation techniques are not sufficient for evaluating deep learning models. These deep learning models should be evaluated on validation dataset for unbiased evaluation.

## Funding Source

The current work has received grant from the Department of Bio-Technology (DBT), Govt of India, India.

## Conflict of interest

The authors declare no competing financial and non-financial interests.

## Authors’ contributions

SP and GPSR collected and processed the datasets. SP, KP, HS and GPSR implemented the algorithms and SP developed the prediction models. SP, AD and GPSR analysed the results. SP created the back-end of the web server the front-end user interface. SP, AD and GPSR penned the manuscript. GPSR conceived and coordinated the project. All authors have read and approved the final manuscript.

## Acknowledgements

Authors are thankful to the Department of Bio-Technology (DBT) and Department of Science and Technology (DST-INSPIRE) for fellowships and the financial support and Department of Computational Biology, IIITD New Delhi for infrastructure and facilities.

## Data Availability Statement

All the datasets generated in this study are available at “Pprint2” web server https://webs.iiitd.edu.in/raghava/pprint2/dataset.php.

## Notes

https://webs.iiitd.edu.in/raghava/pprint2/

### Competing Interest Statement

The authors have declared no competing interest.

